## Supplementary information for "The Immune Factors Driving DNA Methylation Variation in Human Blood"

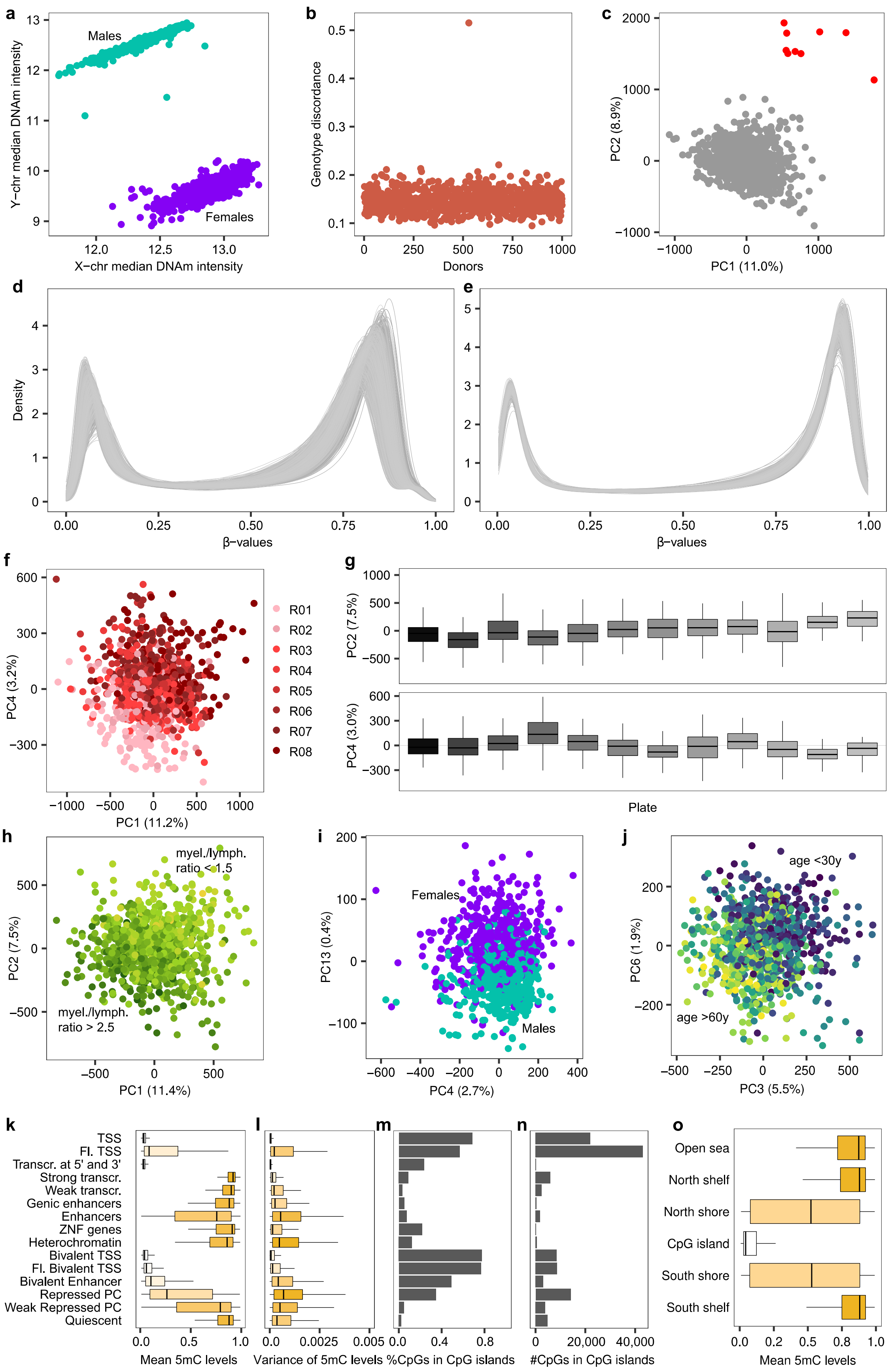
a
Males
Females
Y−chr median DNAm intensity
X−chr median DNAm intensity
b
Genotype discordance
Donors
c
PC2 (8.9%)
PC1 (11.0%)
d
Density
β−values
e
β−values
f
R01
R02
R03
R04
R05
R06
R07
R08
PC4 (3.2%)
PC1 (11.2%)
g
PC2 (7.5%)
PC4 (3.0%)
Plate
h
myel./lymph. ratio < 1.5
myel./lymph. ratio > 2.5
PC2 (7.5%)
PC1 (11.4%)
i
Females
Males
PC13 (0.4%)
PC4 (2.7%)
j
age <30y
age >60y
PC6 (1.9%)
PC3 (5.5%)
k
TSS
Fl. TSS
Transcr. at 5' and 3'
Strong transcr.
Weak transcr.
Genic enhancers
Enhancers
ZNF genes
Heterochromatin
Bivalent TSS
Fl. Bivalent TSS
Bivalent Enhancer
Repressed PC
Weak Repressed PC
Quiescent
Mean 5mC levels
l
Variance of 5mC levels
m
%CpGs in CpG islands
n
#CpGs in CpG islands
o
Open sea
North shelf
North shore
CpG island
South shore
South shelf
Mean 5mC levels

**Supplementary Fig. 1. Processing of the Infinium MethylationEPIC Array Data.**
(**a**) Median DNA methylation intensity on sex chromosomes, in $n$ = 978 Milieu Intérieur donors. Turquoise and violet colors denote donors who declared being men and women, respectively. (**b**) Genotype concordance between DNA methylation and SNP array data, using 59 control SNPs in $n$ = 978 donors. (**c**) Principal component analysis (PCA) of 5mC levels at $n$ = 863,586 CpG sites in $n$ = 978 donors. The red color indicates excluded outliers. (**d**) Epigenome-wide distributions of β values in $n$ = 968 donors, before "noob" background correction. (**e**) Epigenome-wide distributions of β values in $n$ = 968 donors, after "noob" background correction. (**f**) PCA of 5mC levels corrected for background effects at $n$ = 863,906 CpG sites in $n$ = 968 donors. Colors indicate sentrix positions. (**g**) Distribution of principal components (PC) across sample plates, obtained by PCA of 5mC levels corrected for background and sentrix position effects, at $n$ = 863,906 CpG sites in $n$ = 968 donors. (**h-j**) PCA of 5mC levels corrected for background, sentrix position and sample plate effects, at 644,517 CpG sites in $n$ = 884 donors. Colors indicate (**h**) the log-ratio of the proportions of myeloid vs. lymphoid lineages in blood, (**i**) declared sex and (**j**) age. (**k**) Distributions of 5mC levels at $n$ = 644,517 CpG sites averaged over $n$ = 884 donors, across 15 chromatin states. (**l**) Distributions of the variance of 5mC levels at $n$ = 644,517 CpG sites among $n$ = 884 donors, across 15 chromatin states. (**m**) Proportion of CpG sites in CpG islands, across 15 chromatin states. (**n**) Number of CpG sites in CpG islands, across 15 chromatin states. (**o**) Distributions of 5mC levels at $n$ = 644,517 CpG sites averaged over $n$ = 884 donors, according to distance to CpG islands. (**c, f-j**) The proportion of variance explained by principal components (PC) is indicated in brackets. (**g, k, l, o**) The line, box and whiskers indicate the median, interquartile range and 1.5× the interquartile range, respectively.

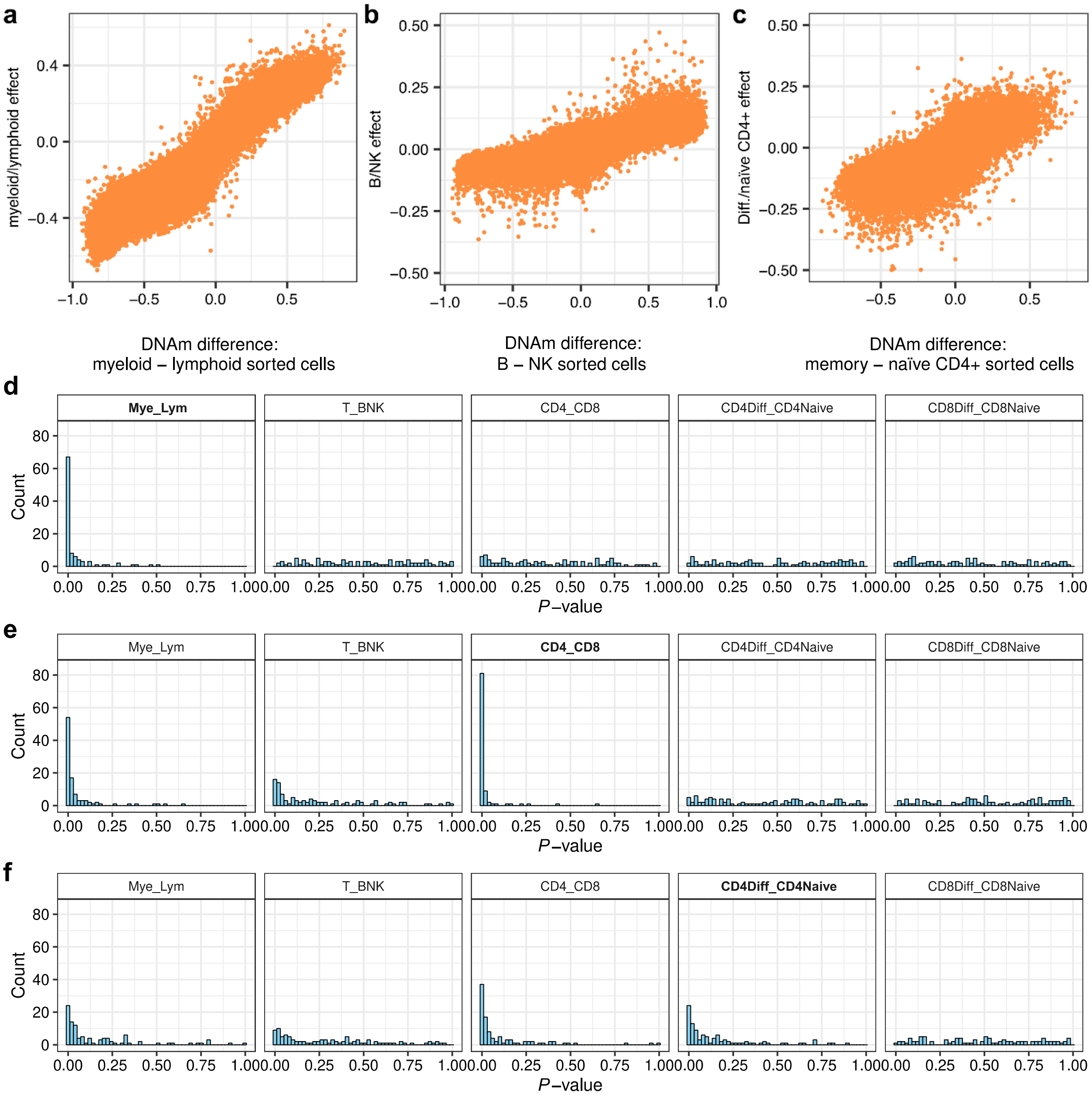


**Supplementary Fig. 2. Validation of the Models Estimating the Effects of Cellular Composition on DNA Methylation**

(**a**) Estimated effects of the myeloid/lymphoid log-ratio on DNA methylation, against the measured differences in DNA methylation between sorted cells from the myeloid and lymphoid lineages[28]. (**b**) Estimated effects of the B cell/NK cell log-ratio on DNA methylation, against the measured differences in DNA methylation between sorted B and NK cells[28]. (**c**) Estimated effects of the differentiated/naïve $CD4^+$ T cell log-ratio on DNA methylation, against the measured differences in DNA methylation between sorted, differentiated and naïve $CD4^+$ T cells[28]. (**d**) *P*-value histograms for simulations where myeloid and lymphoid cells have differentially distributed 5mC levels. (**e**) *P*-value histograms for simulations where $CD4^+$ and

CD8$^+$ T cells have differentially distributed 5mC levels. (**f**) *P*-value histograms for simulations where differentiated and naïve CD4$^+$ T cells have differentially distributed 5mC levels.

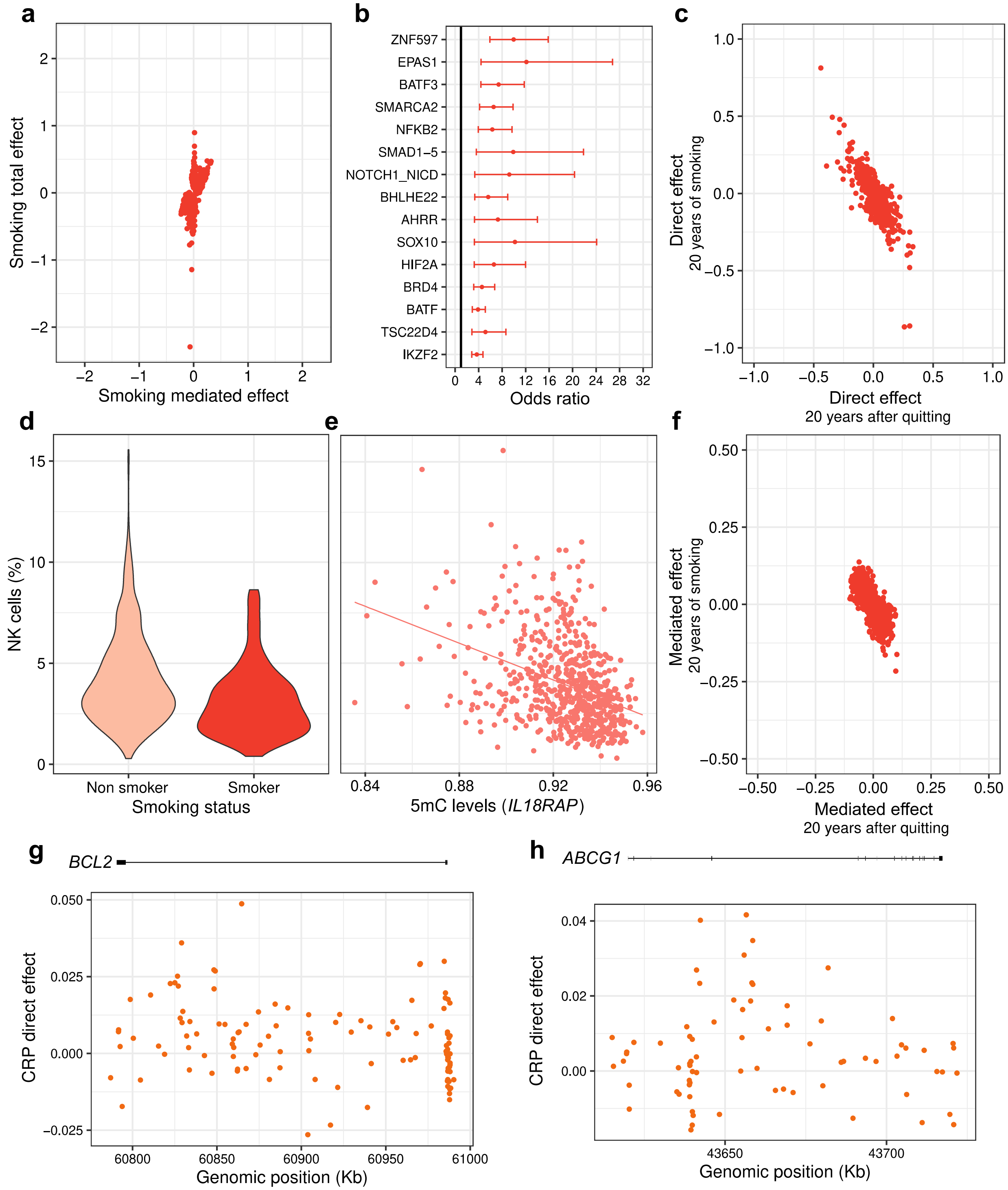


**Supplementary Fig. 3. The Effects of Environmental Exposures on the Blood DNA Methylome of Adults**

(**a**) Total effects against cell-composition-mediated effects of smoking on 5mC levels. Only CpG sites with a significant total and/or cell-composition-mediated effect are shown. (**b**) Enrichment of CpG sites with significant direct, negative effects of smoking in binding sites for TFs. The point and error bars indicate the odds-ratio and 95% CI. CIs were estimated by the Fisher's exact method. The 15 most enriched TFs are shown, out of 1,165 tested TFs. (**c**) Direct effects of 20 years of cigarette smoking against direct effects of 20 years without cigarette smoking on 5mC levels. Only CpG sites with a significant direct effect of smoking are shown. (**d**) Distributions of the proportion of NK cells in non-smokers and smokers. (**e**) 5mC levels at the *IL18RAP* locus against the proportion of NK cells. 5mC levels are given in the β value scale. (**f**) Cell-composition-mediated effects of 20 years of cigarette smoking against cell-composition-mediated effects of 20 years without cigarette smoking on 5mC levels. Only CpG sites with a significant cell-composition-mediated effect of smoking are shown. (**g**) Genomic distribution of direct effects of C-reactive protein (CRP) levels at the *BCL2* locus. (**h**) Genomic distribution of direct effects of CRP levels at the *ABCG1* locus. (**a, c, f-h**) Effect sizes are given in the M value scale. Statistics were computed based on a sample size of $n$ = 884 and for 644,517 CpG sites.

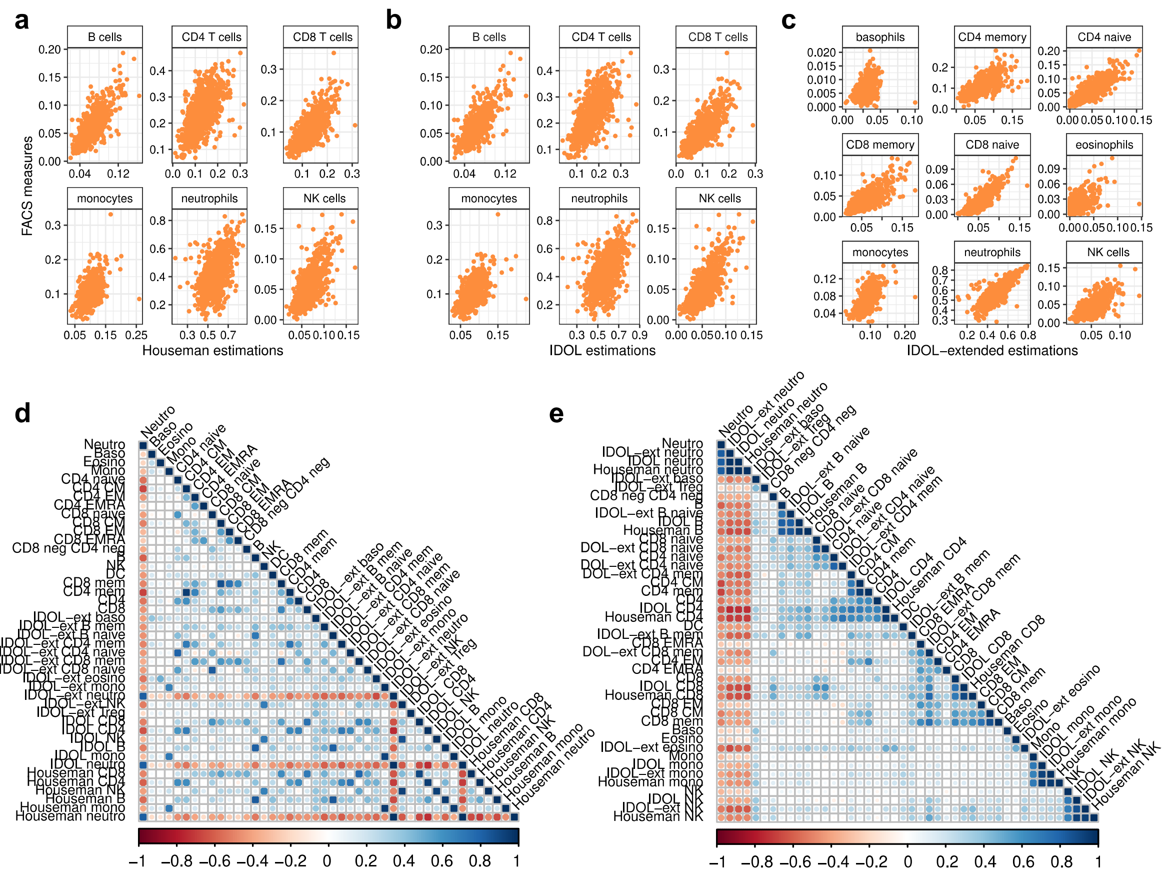


**Supplementary Fig. 4. Measured against Estimated Blood Cell Proportions**
(**a**) Proportions of six blood cell subsets measured by flow cytometry in *n* = 884 Milieu Intérieur donors, against proportions estimated by the Houseman *et al.*'s cell mixture deconvolution method. (**b**) Proportions of six blood cell subsets measured by flow cytometry in *n* = 884 Milieu Intérieur donors, against proportions estimated by the IDOL cell mixture deconvolution method. (**c**) Proportions of nine blood cell subsets measured by flow cytometry in *n* = 884 Milieu Intérieur donors, against proportions estimated by the EPIC IDOL-Ext cell mixture deconvolution method. (**d**) Matrix of pairwise correlation coefficients between measured and estimated blood cell proportions. Cell proportions are ordered according to data type. (**e**) Matrix of pairwise correlation coefficients between measured and estimated blood cell proportions. Cell proportions are ordered according to correlation coefficients.

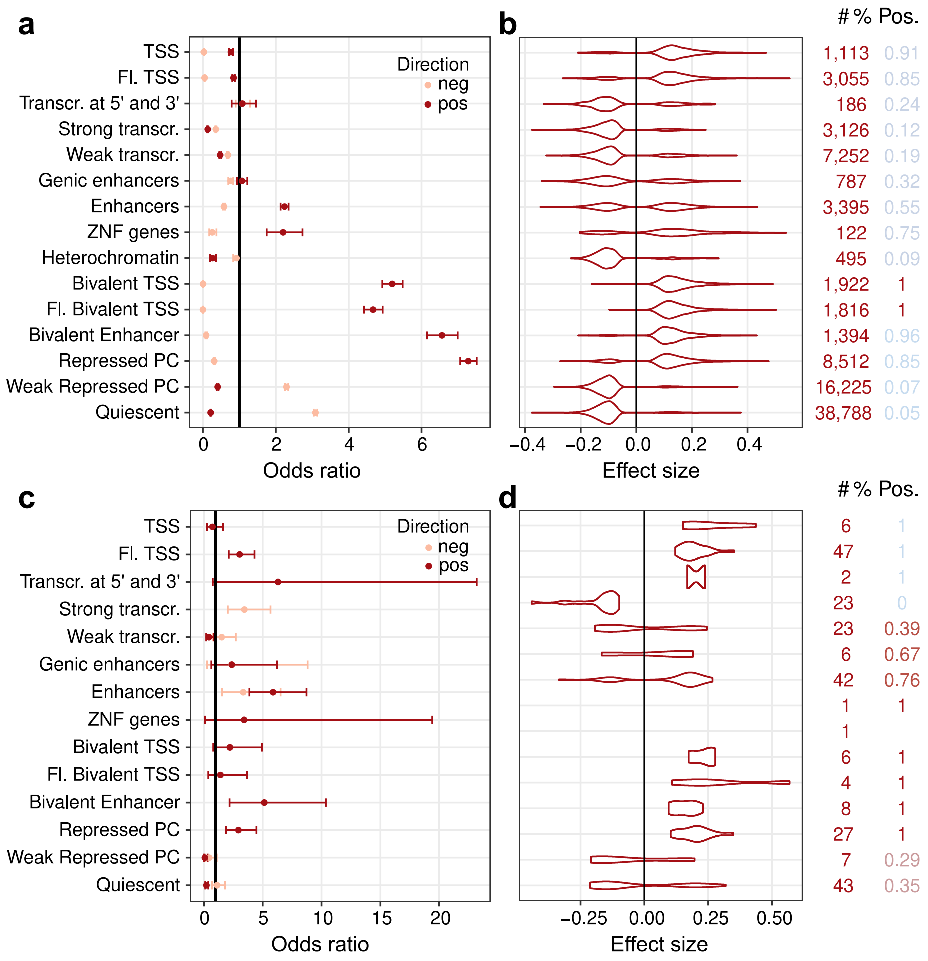


**Supplementary Fig. 5. The Effects of CMV infection on the Blood DNA Methylome of Adults**

(**a**) Enrichment in CpG sites with significant cell-composition-mediated effects of CMV infection, across 15 chromatin states. (**b**) Distributions of significant cell-composition-mediated effects of CMV infection, across 15 chromatin states. (**c**) Enrichment in CpG sites with significant direct effects of CMV infection, across 15 chromatin states. (**d**) Distributions of significant direct effects of CMV infection, across 15 chromatin states. (**a, c**) The point and error bars indicate the odds-ratio and 95% CI. CIs were estimated by the Fisher's exact method. Statistics were computed based on a sample size of $n$ = 884 and for 644,517 CpG sites.

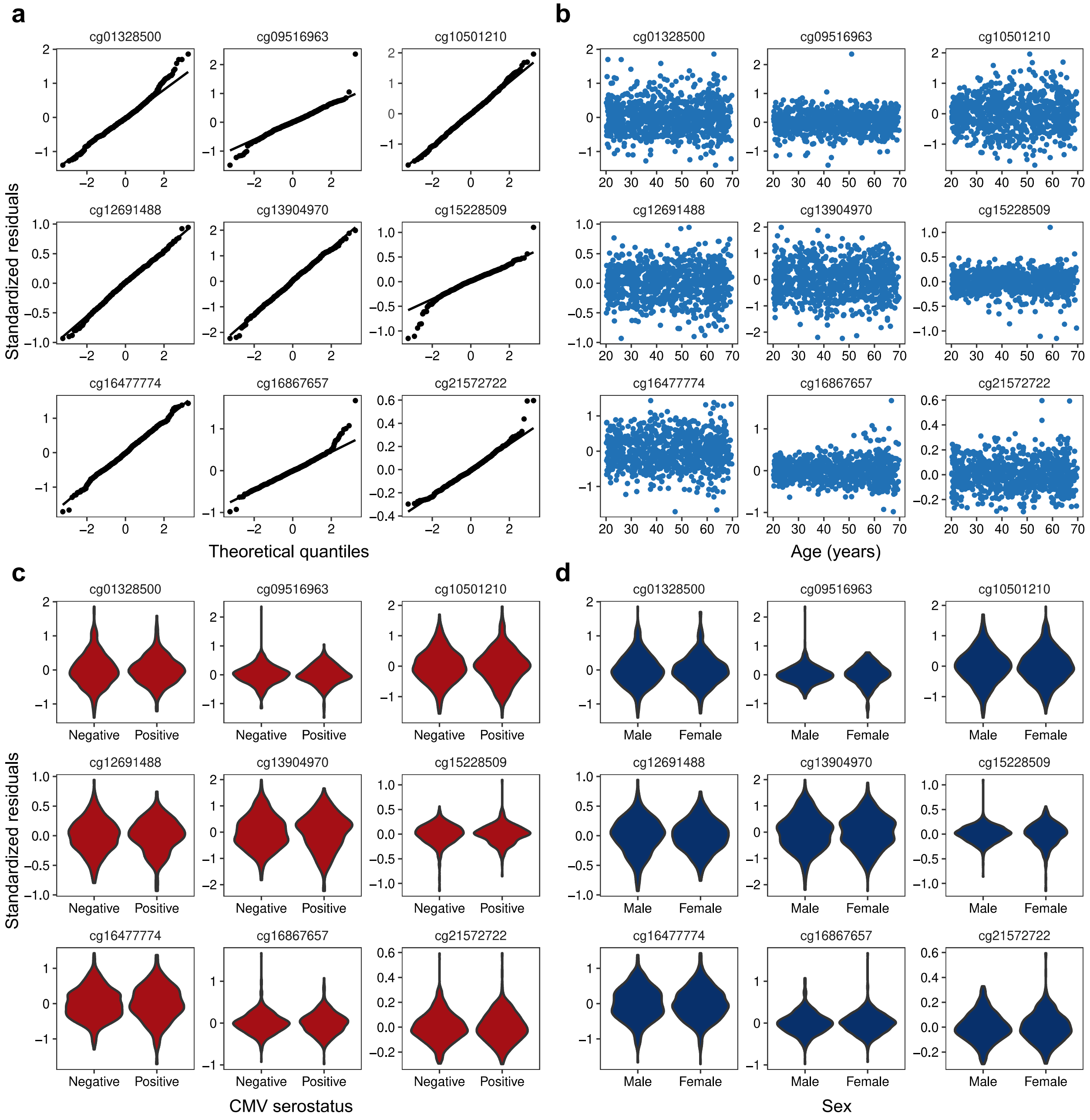


**Supplementary Fig. 6. Diagnostics for Regression Models of DNA Methylation Levels Associated with Age, Sex and CMV infection**

(**a**) Q-Q plots of residuals for multiple regression models of DNA methylation levels at the CpG sites the most associated with age, sex and CMV infection. (**b**) Residuals against age. (**c**) Residuals against CMV serostatus. (**d**) Residuals against sex. Statistics were computed based on a sample size of $n = 884$ and for 644,517 CpG sites.

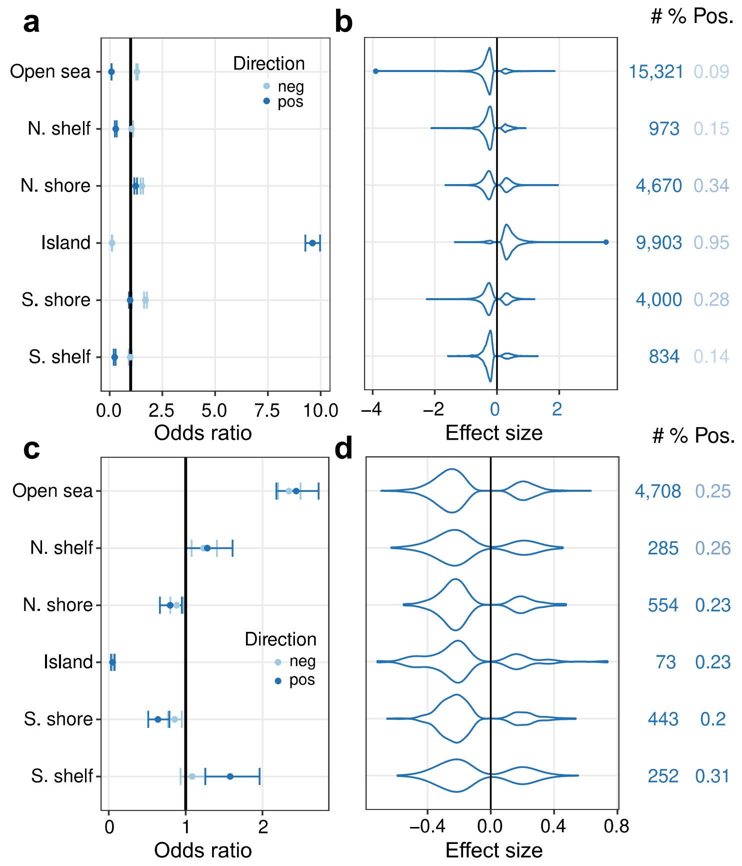


**Supplementary Fig. 7. The Effects of Age on the Blood DNA Methylome of Adults**
(**a**) Enrichment in CpG sites with significant direct effects of age, according to distance to CpG islands. (**b**) Distributions of significant direct effects of age, according to distance to CpG islands. (**c**) Enrichment in CpG sites with significant cell-composition-mediated effects of age, according to distance to CpG islands. (**d**) Distributions of significant cell-composition-mediated effects of age, according to distance to CpG islands (**a-d**) Effect sizes are given in the M value scale. (**a, c**) The point and error bars indicate the odds-ratio and 95% CI. CIs were estimated by the Fisher's exact method. (**b, d**) Numbers on the right indicate the number of associated CpG sites and proportion of positive effects. Statistics were computed based on a sample size of $n$ = 884 and for 644,517 CpG sites.

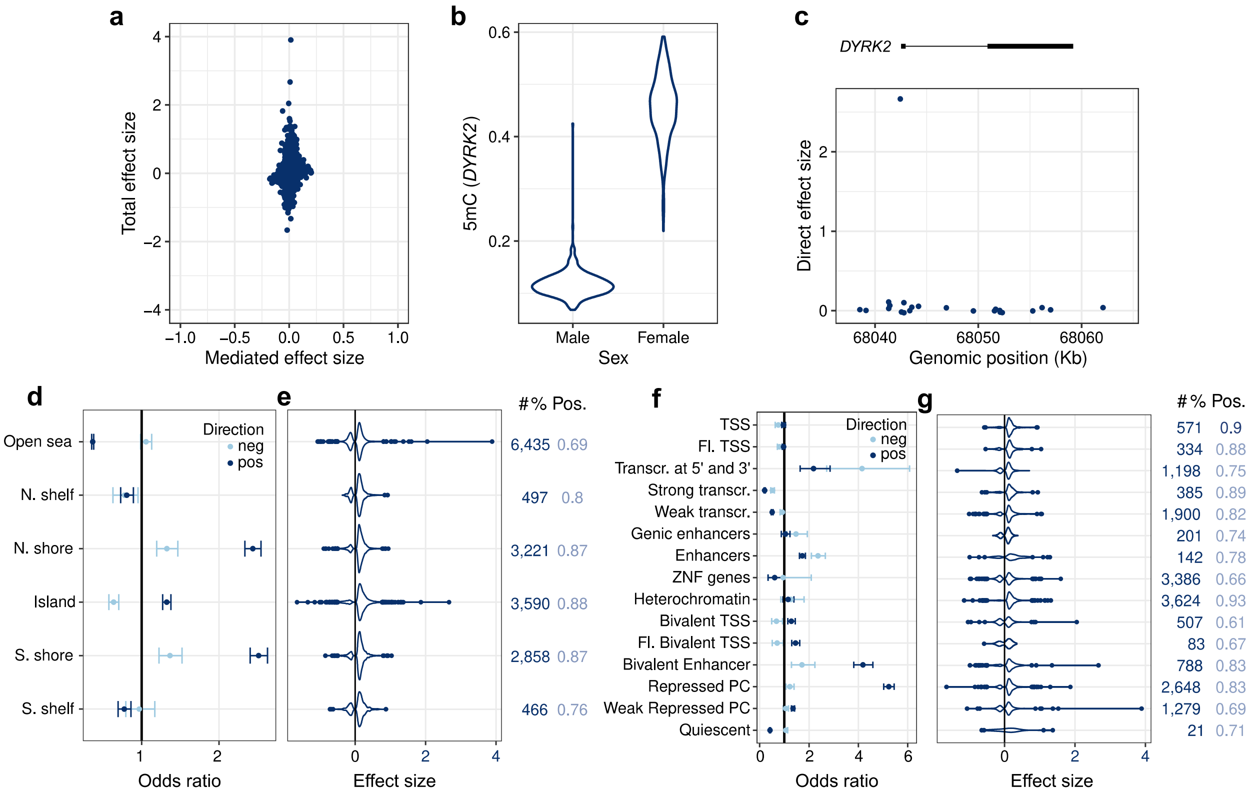


**Supplementary Fig. 8. The Effects of Sex on the Blood DNA Methylome of Adults**
(**a**) Total effects against cell-composition-mediated effects of sex on 5mC levels. Only CpG sites with a significant total and/or cell-composition-mediated effect are shown. (**b**) Distributions of 5mC levels at the *DYRK2* locus in women and men. 5mC levels are given in the β value scale. (**c**) Genomic distribution of direct sex effects at the *DYRK2* locus. (**d**) Enrichment in CpG sites with significant direct effects of sex, according to distance to CpG islands. (**e**) Distributions of significant direct effects of sex, according to distance to CpG islands. (**f**) Enrichment in CpG sites with significant direct effects of sex, across 15 chromatin states. (**g**) Distributions of significant direct effects of sex, across 15 chromatin states. (**d, f**) The point and error bars indicate the odds-ratio and 95% CI. CIs were estimated by the Fisher's exact method. (**e, g**) Numbers on the right indicate the number of associated CpG sites and proportion of positive effects. (**a, c, e, g**) Effect sizes are given in the M value scale. Statistics were computed based on a sample size of $n$ = 884 and for 644,517 CpG sites.

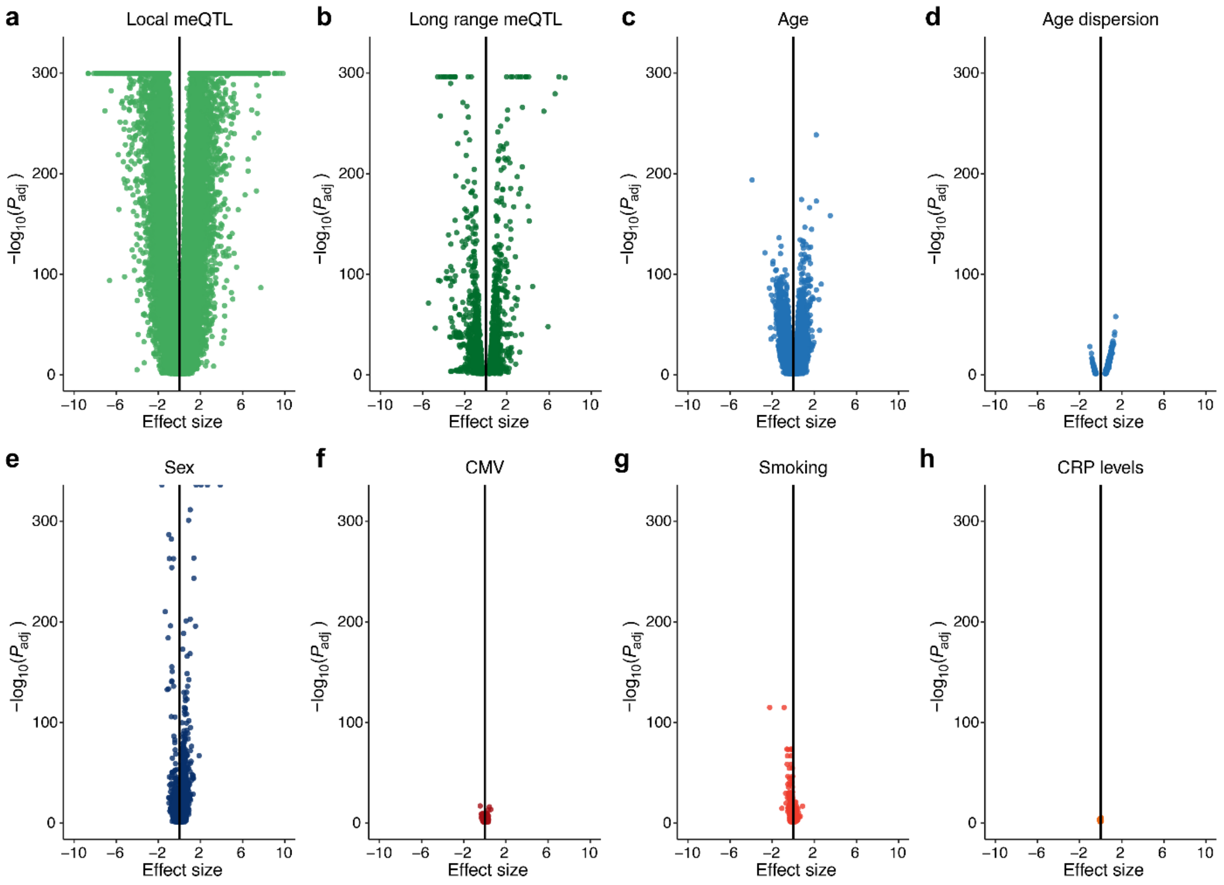


**Supplementary Fig. 9. Volcano Plots of Genetic, Intrinsic and Exposure Effects on the Blood DNA Methylome of Adults.** Effect sizes and *P*-values were estimated from linear regression models of DNA methylation levels with (**a**) local meQTL variants, (**b**) remote meQTL variants, (**c**) age, (**d**) age dispersion, (**e**) sex, (**f**) CMV serostatus, (**g**) smoking status and (**h**) log-transformed CRP levels as predictors (Methods). Scales are identical across plots, to facilitate comparisons. Effect sizes are given in the M value scale. Statistics were computed based on a sample size of $n = 884$ and for 644,517 CpG sites.

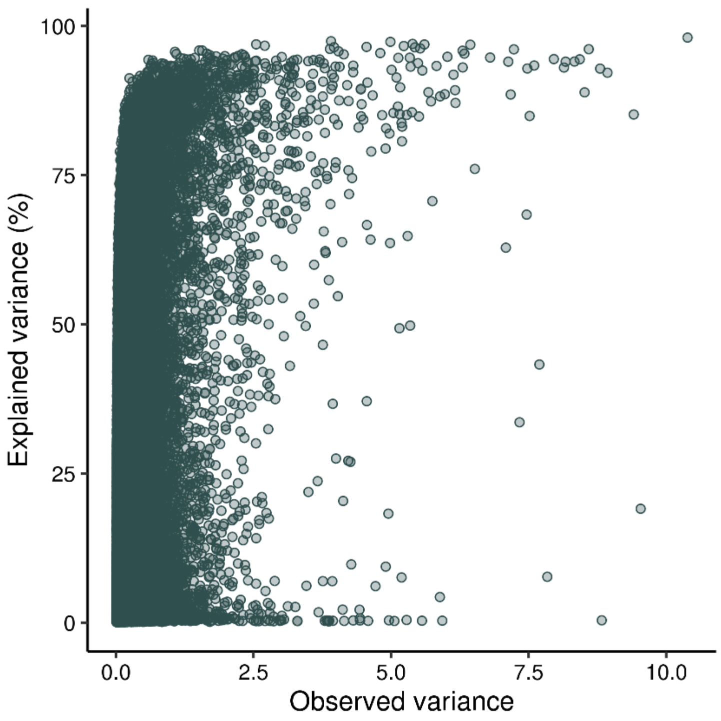


**Supplementary Fig. 10. Observed and Explained Variance of the Blood DNA Methylome of Adults.** Observed variance against the proportion of variance explained by blood cell composition, genetic and intrinsic factors and exposures.

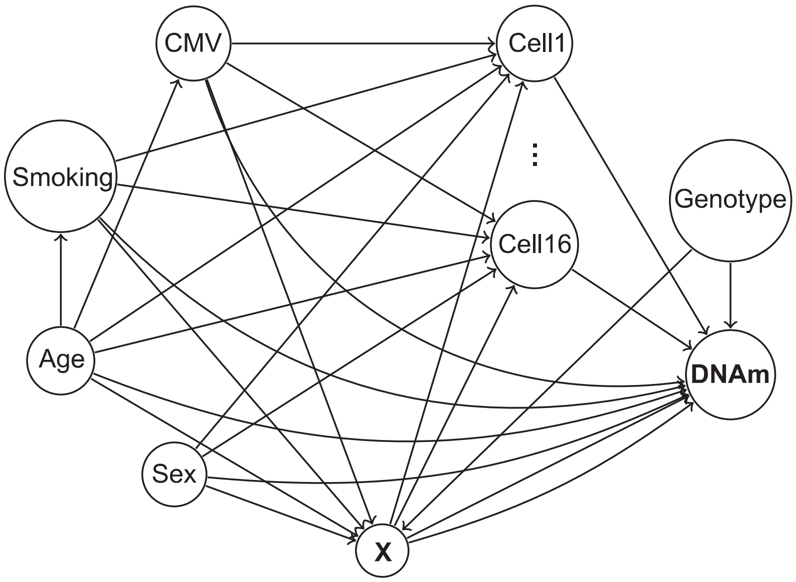


**Supplementary Fig. 11. Causal Relationships Assumed in the Present Study.** Directed acyclic graph representing causal relationships assumed between DNA methylation, a tentative environmental exposure X, genetic, intrinsic and environmental factors, and blood cell composition ($\mathrm{Cell}_i, i = 1, \cdots, 16$).

### Supplementary Notes

**Population Variation in DNA Methylation Across the Genome.** To identify the genomic regions where DNA methylation is the least and most variable, we estimated the average and variance of DNA methylation among Milieu Intérieur donors, across different well-characterized chromatin states, using peripheral blood mononuclear cells (PBMCs) as a reference[1]. We found that CpG sites in transcription start sites (TSS) are typically unmethylated and exhibit the lowest population variance in 5mC levels (Supplementary Fig. 1k,l), suggesting that epigenetic constraints are the strongest in promoters, whereas actively transcribed gene bodies are highly methylated and also show low population variance. 5mC levels in enhancers and Polycomb-repressed regions are the most variable (Supplementary Fig. 1l), suggesting that DNA methylation in these regions are preferentially affected by genetic, intrinsic or environmental factors, or cellular heterogeneity. Expectedly, we observed that CpG sites within and outside of CpG islands are mainly demethylated and methylated, respectively (Supplementary Fig. 1o). These results indicate that 5mC measurements from our cohort reproduce the known properties of DNA methylation and show high levels of variation across the epigenome and among individuals.

**Simulation-based validation of associations between immune cell composition and 5mC levels.** We performed a simulation study to validate the models used to estimate the association of blood cell composition with 5mC levels. We simulated proportions of 16 cell-types from a Dirichlet distribution, with shape parameters $\alpha_c$, with $c = 1, \ldots, 16$, sampled from a log-normal distribution, $\log \alpha_c \sim \mathcal{N}(0,1)$. We ordered shape parameters to make the order of the simulated average proportions for the 16 cell-types correspond to the order of average proportions of the cell-types in the observed samples. For a particular CpG site, we simulated 5mC levels differently in two groups of immune cells, sampled from two different beta distributions with the "default" group having parameters $a = 10$ and $b = 1$, and the "alternative" group having parameters $a = 5$ and $b = 1$, respectively. Let $\boldsymbol{\theta}$ be simulated proportions and $\boldsymbol{\beta}$ simulated 5mC levels for the16 cell-types for one individual. 5mC levels at a given CpG site was the computed by $\boldsymbol{\theta}^t\boldsymbol{\beta}$**.** We simulated 5mC levels for 1,000 individuals at100 CpG sites. For each CpG site, we then fitted a regression model including log-ratios of groups of immune cell proportions as predictors (Methods).We considered three different scenarios with different groups of cell-types in the “alternative” group. For the first scenario, all myeloid cell-types were simulated to belong to the alternative group. As expected, we found a strong effect of the myeloid-lymphoid log-ratio (referred to as a balance) and no effect of other balances (Supplementary Fig. 2d). For the second scenario, $CD4^+$ T cells were simulated to be in the alternative group. In this case we found an effect of the balances encoding relative DNA methylation differences of myeloid and lymphoid cells, T cells with NK and B cells, and $CD4^+$ T cells with $CD8^+$ T cells (Supplementary Fig. 2e), which is expected since $CD4^+$ T cells are included in the lymphoid and T cell groups. Finally, in the third scenario we simulated differentiated $CD4^+$ T cells (CM, EM or EMRA $CD4^+$ T cells) to be in the alternative group. As expected for this scenario, we found an effect of the balances encoding differences between myeloid and lymphoid cells, T cells with NK and B cells, CD4+ T cells and CD8+ T cells, and differentiated $CD4^+$ T cells with naïve $CD4^+$ T cells, but not for differentiated $CD8^+$ T cells and naïve $CD8^+$ T cells (Supplementary Fig. 2f). Together, these results indicate that models including balances of immune cell proportions as predictors can accurately estimate differences in DNA methylation between different hierarchical groups of cell-types.

**Strong Effects of Smoking are Independent of Blood Cell Composition.** Although active smoking is known to elicit reproducible changes in DNA methylation[2,3], we previously showed that tobacco consumption also has a broad effect on blood immune cell subsets, particularly B and NK cells[4], suggesting possible mediation by cellular composition. When adjusting for the 16 immune cell proportions, we found that smoking directly alters 5mC levels at 428 CpG sites (~0.07% of CpG sites; Fig. 1b and Supplementary Table 1), 70.8% of which show a decrease in 5mC levels (Supplementary Fig. 3a). Smokers show strongly decreased 5mC levels in the introns of the dioxin receptor repressor gene *AHRR* (β value scale 95% CI: [-23%, -20%], $P_{adj} = 1.2\times10^{-115}$), the second exon of *F2RL3* (CI: [-9.9%, -8.1%], $P_{adj} = 1.8\times10^{-67}$) and the first intron of *RARA* (CI: [-10.4%, -8.4%], $P_{adj} = 3.0\times10^{-59}$), in agreement with previous studies[2,3]. No clear differences in the distribution of effect sizes were observed across chromatin states. CpG sites that are demethylated in smokers are significantly enriched in binding sites for the hypoxia-related TFs EPAS1 and HIF2A (OR > 6.0, $P < 2.0\times10^{-6}$) and AHRR (OR = 7.3, CI: [3.3, 14.0], $P = 7.0\times10^{-6}$; Supplementary Fig. 3b and Supplementary Table 3), implying that AHRR up-regulation in smokers elicits decreased 5mC levels at AHRR binding sites. To determine if such direct effects are reversible, we compared, for all smoking-associated CpG sites, the changes in 5mC levels with years since last smoke for past smokers, to the changes with years of smoking for active smokers. We found a strong negative correlation between effect sizes ($R$ = -0.82; slope = -1.08; Supplementary Fig. 3c), supporting the reversibility of the direct effects of smoking on DNA methylation[2,3].

We estimated that 5mC levels are significantly altered by smoking due to changes in cell composition at ~0.58% CpG sites ($n$ = 3,713 CpG sites with a mediated effect; $P_{adj} < 0.05$; Supplementary Fig. 3a). Among the most strongly affected CpG sites, we found *IL18RAP* (CI: [0.56%, 0.92%], $P_{adj} = 6.80\times10^{-11}$), a subunit of the receptor for IL18 that is differentially expressed by NK cells[5]. We observed that active smoking induces a reduction in the proportion of NK cells ($P = 2.91\times10^{-8}$), which is in turn associated with lower 5mC levels at *IL18RAP* ($P = 5.83\times10^{-32}$; Supplementary Fig. 3d, e), in line with an effect of smoking mediated by NK cells. Of note, mediated effects of smoking on 5mC levels are also reversible, to a degree similar to that of direct effects ($R$ = -0.78; slope=-0.99, Supplementary Fig. 3f). Importantly, only three out of the 428 CpG sites with a direct effect of smoking status also have a significant mediation effect, and, on average, mediated effects of smoking are much weaker than direct effects (Supplementary Fig. 3a). Furthermore, we found no evidence, using an interaction model, that smoking effects on DNA methylation depend on the proportion of myeloid cells, in contrast with previous studies, including at the *GPR15* gene[6,7]. Collectively, these findings indicate that the largest effects of cigarette smoking on the blood DNA methylome are reversible and independent of blood cell composition.

**Effects of Other Environmental Exposures on the Adult DNA Methylome.** Another exposure that we identified as affecting DNA methylation variation is circulating levels of C-reactive protein (CRP), a marker of chronic, low-grade inflammation in healthy adults. Associations between CRP levels and hundreds of 5mC marks have been detected[8], but the strong relationship of CRP levels with the immune system[9] and genetic variation[10] suggests that that these factors could confound associations. Specifically, changes in blood cell composition may cause changes in CRP levels, and this could induce spurious associations, instead of mediated effects, at CpG sites associated with immune cell proportions.

We found a significant total effect of CRP levels on DNA methylation at 53 CpG sites ($P_{adj}$ < 0.05; Table 2), a figure that, when adjusting for cellular composition, decreases to 39, of which 77% ($n$ = 30) show decreased 5mC levels with increased CRP levels (Fig. 1b and Supplementary Table 1). We detected a CpG site within an enhancer nearby *BCL2,* a key regulator of apoptosis and inflammation[11], where 5mC levels increase with increasing CRP levels (β value scale 95% CI: [0.6%, 1.1%], $P_{adj} = 4.9\times10^{-6}$; Supplementary Fig. 3g). Another example is a CpG site within an enhancer nearby *ABCG1* (CI: [0.5%, 0.9%], $P_{adj} = 6.4\times10^{-6}$; Supplementary Fig. 3h). In our cohort, 5mC levels at the same site are also associated with triglyceride (CI: [1.2%, 2.1%], $P_{adj} = 1.2\times10^{-7}$) and HDL (CI: [-3.7%, -1.8%], $P_{adj} = 2.5\times10^{-3}$) levels. The associations are retained in a model including CRP, HDL and triglyceride levels, indicating that they affect *ABCG1* 5mC levels independently. CRP is known to inhibit cellular cholesterol efflux by downregulating *ABCG1* mRNA levels, which are impaired in patients with type 2 diabetes, obesity, and hypertension[12]. These results suggest that several associations between CRP levels and DNA methylation are not due to changes in blood cell composition, and generate new hypotheses on the epigenetic mechanisms relating subclinical inflammation to metabolic conditions.

Besides CMV infection, smoking status and chronic inflammation, we found limited evidence of a direct effect of exposures on the DNA methylome of healthy adults (Supplementary Table 1). In total, we found 36 significant direct associations between the remaining non-heritable factors and 5mC levels, 12 of which relate *ABCG1*, *DHCR24* and *CPT1A* genes with lipid-related traits and BMI, as previously reported[13]. We also found a significant association between log uric acid levels and 5mC levels at the *SLC2A9* gene (cg00071950; $P_{adj}$ = 0.0034), which is no longer significant when adjusting on the local meQTL SNP ($P_{adj}$ = 1.0), illustrating how DNA sequence variation can confound EWAS results. Nutritional habits, assessed based on 20 dietary frequency variables, have no detectable effects on the blood DNA methylome, except the frequency of raw fruit consumption at *GLI2* (95% CI: [-2.5%, -1.2%], $P_{adj}$ = 0.0075). Of note, we did not replicate previously reported associations between DNA methylation and serum IgE levels[14,15] and did not detect any association with current socio-economic status. Collectively, these results indicate that environmental exposures related to upbringing, socio-economic status, nutrition or vaccination do not induce strong changes of the blood DNA methylome in our cohort of healthy adults.
